# Loss of flavin adenine dinucleotide (FAD) impairs sperm function and male reproductive advantage in *C. elegans*

**DOI:** 10.1101/456863

**Authors:** Chia-An Yen, Dana L. Ruter, Christian D. Turner, Shanshan Pang, Sean P. Curran

**Affiliations:** Leonard Davis School of Gerontology, University of Southern California, Los Angeles, CA 90089; Department of Molecular and Computation Biology, Dornsife College of Letters, Arts, and Sciences, University of Southern California, Los Angeles, CA 90089; School of Life Sciences, Chongqing University, Chongqing 401331, China; Norris Comprehensive Cancer Center, Keck School of Medicine, University of Southern California, Los Angeles, CA 90089

**Keywords:** spermatogenesis, mitochondria, germ cells, reproduction, proline catabolism, alh-6/ALDH4A1, C. elegans, senescence, aging, male-specific, FAD, flavin cofactor, riboflavin

## Abstract

Exposure to environmental stress is clinically established to influence male reproductive health, but the impact of normal cellular metabolism on sperm quality is less well-defined. Here we show that impaired mitochondrial proline catabolism, reduces energy-storing flavin adenine dinucleotide (FAD) levels, alters mitochondrial dynamics toward fusion, and leads to age-related loss of sperm quality (size and activity), which reduces competitive fitness. Loss of the 1-pyrroline-5-carboxylate dehydrogenase enzyme *alh-6* that catalyzes the second step in mitochondrial proline catabolism, leads to premature male reproductive senescence. Reducing the expression of the proline catabolism enzyme *alh-6* or FAD biosynthesis pathway genes in the germline is sufficient to recapitulate the sperm-related phenotypes observed in *alh-6* loss-of-function mutants. These sperm-specific defects are suppressed by feeding diets that restore FAD levels. Our results define a cell autonomous role for mitochondrial proline catabolism and FAD homeostasis on sperm function and specify strategies to pharmacologically reverse these defects.

## INTRODUCTION

As individuals wait longer to have families, reproductive senescence has become an increasingly prudent topic (Lemaitre & Gaillard; Mills et al.). Decline in oocyte quality is well-documented with age and can result in fertility issues when older couples try to conceive (Baird et al.). Furthermore, pregnancies at an older age pose risks for higher incidences of birth defects and miscarriages. In humans, female reproduction ceases at an average age of 41-60, with the onset of menopause (Treloar). The *Caenorhabditis elegans* "wild type" is hermaphroditic and self-fertilizing; however, they are capable of making and maintaining Mendelian ratios of male (sperm-only) animals in their populations. Like humans, *C. elegans* experience a decline in fecundity with age by halting oocyte production at roughly one-third of their lifespan (Kadandale & Singson). In addition, regulators of reproductive aging, such as insulin/IGF-1 and *sma-*2/TGF-β signaling, are conserved regulators of reproductive aging from worms to humans (Luo, Kleemann, Ashraf, Shaw, & Murphy). While the majority of studies in reproductive senescence have focused on maternal effects, male factors contribute to a large portion of fertility complications with increasing evidence of an inverse relationship between paternal age and sperm health (Lemaitre & Gaillard). In fact, studies in mammals have shown an age-related decline in sperm quality with increased incidences of DNA damage, reduced motility, abnormal morphology, and decreased semen volume (Cocuzza et al.; Kidd, Eskenazi, & Wyrobek; Ozkosem, Feinstein, Fisher, & O’Flaherty).

Flavin adenine dinucleotide (FAD) is an important cofactor that participates in enzymatic redox reactions that are used in cellular metabolism and homeostasis. FAD is synthesized from riboflavin by the concerted actions of FAD synthetase and riboflavin kinase. Like humans, *C. elegans* cannot synthesize riboflavin, and therefore requires dietary intake (Braeckman, Houthoofd, & Vanfleteren). Disruption of flavin homeostasis in humans and animal models has been associated with several diseases, including: cardiovascular diseases, cancer, anemia, abnormal fetal development, and neuromuscular and neurological disorders (Barile et al.).; however, a link between FAD homeostasis and fertility is undefined.

Several studies have documented fertility defects in *C. elegans* mitochondrial mutants. Mutation in *nuo-1*, a complex I component of the mitochondria respiratory chain, results in reduced brood size caused by impaired germline development (Grad & Lemire). Similarly, *clk-1* mutation affects the timing of egg laying, resulting in reduced brood size (Jonassen, Marbois, Faull, Clarke, & Larsen). Both of these mitochondrial mutations impact fertility, but their role(s) in spermatogenesis are unclear. *alh-6*, the *C. elegans* ortholog of human *ALDH4A1*, is a nuclear-encoded mitochondrial enzyme that functions in the second step of the proline metabolism pathway, converting 1-pyrroline-5-carboxylate (P5C) to glutamate (Adams & Frank). We previously revealed that *alh-6(lax105)* loss-of-function mutants display altered mitochondrial structure in the muscle accompanied by increased level of ROS in adult animals (Pang & Curran). Furthermore, mutation in *alh-6* results in the activation of SKN-1/NRF2 (Pang, Lynn, Lo, Paek, & Curran), an established regulator of oxidative stress response, likely through the accumulation of toxic P5C disrupting mitochondrial homeostasis (Deuschle et al.; Miller et al.; Nomura & Takagi; Pang & Curran; Pang et al.). Interestingly, SKN-1 was recently shown to respond to accumulation of damaged mitochondria by inducing their biogenesis and degradation through autophagy (Palikaras, Lionaki, & Tavernarakis). Here, we identify a genetic pathway for regulating male reproductive decline stemming from perturbation of mitochondrial proline metabolism leading to redox imbalance, cofactor depletion, and altered mitochondria dynamics; all of which play a role in sperm dysfunction.

## RESULTS

### Mutation in mitochondrial *alh-6* results in diet-independent reduction in fertility

Altered mitochondrial structure and activity have been correlated with sperm dysfunction across different species (Amaral, Lourenco, Marques, & Ramalho-Santos; Liau, Gonzalez-Serricchio, Deshommes, Chin, & LaMunyon; Nakada et al.; Ramalho-Santos & Amaral). In addition, proper sperm function requires low levels of ROS (de Lamirande & Gagnon; Kodama, Kuribayashi, & Gagnon; Leclerc, de Lamirande, & Gagnon), although a specific role for endogenous mitochondrial derived ROS is undefined. ALH-6/ALDH4A1, is a nuclear-encoded mitochondrial enzyme that functions in the second step of proline catabolism, converting 1-pyrroline-5-carboxylate (P5C) to glutamate (Figure 1A). We anticipated that mutation of *alh-6* may affect the germline, based on our previous assessment of the premature aging phenotypes in somatic cells in *alh-6* mutants (Pang & Curran). Using a UV-integrated *alh-6::gfp* strain under its endogenous promoter, we saw that ALH-6 localizes to the mitochondria in the germline of both hermaphrodites (Figure S1A-B) and males (Figure S1C-D). We then assessed progeny output of *alh-6(lax105)* hermaphrodites fed the standard OP50/*E. coli* B strain diet and found a reduction in self-fertility brood size (−12.9%) (Figure 1B). Since the somatic phenotypes of *alh-6(lax105)* mutants are known to be diet-dependent (Pang & Curran; Pang et al.), we examined self-fertility of animals fed the HT115/*E. coli* K-12 strain diet to determine if the reduced reproductive output is also dependent on the type of bacterial diet ingested. Surprisingly, we found that the self-fertility of *alh-6* animals was markedly reduced (−20.7%), when animals were fed the HT115 diet (Figure 1C). *alh-6* mutants have similar timing in their progeny output as compared to wild type animals on both diets (Figures S2A-B). Since *alh-6* mutants display normal development and reproductive timing, the progeny deficit is not a result of an attenuated reproductive span which reveals the differential impact of *alh-6* loss in the soma (diet-dependent) (Pang & Curran) and the germline (diet-independent)

**Figure 1.**
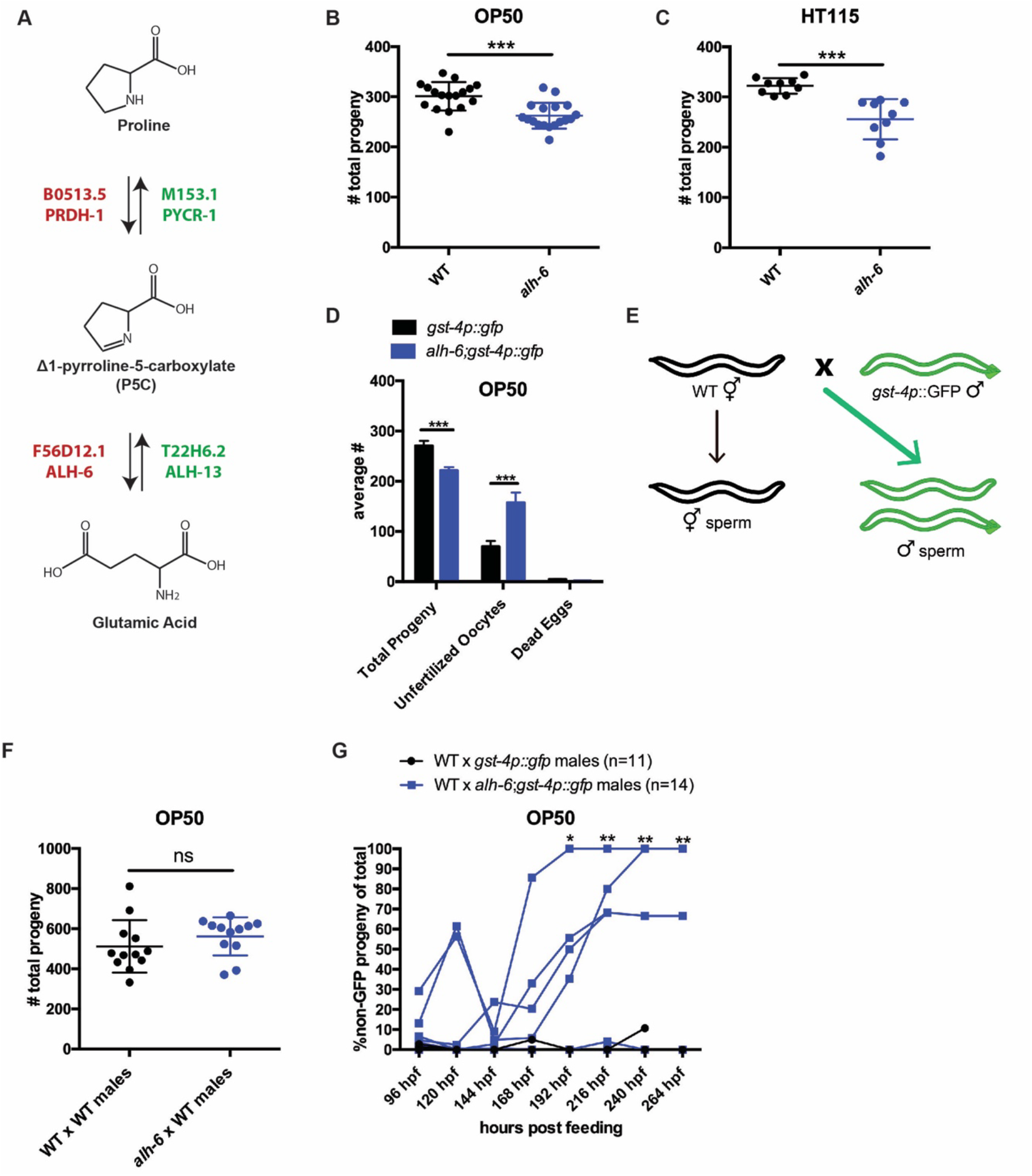
*alh-6* fertility defects are sperm-specific. (A) Proline catabolism pathway. (B-C) *alh-6* hermaphrodites have reduced brood size when fed OP50 (B) or HT115 (C) diets. (D) *alh-6* hermaphrodites produce increased unfertilized oocytes, but few dead embryos. (E) Mated reproductive assay scheme utilizes males to maximize reproductive output (as in F) and can exploit males harboring GFP to differentiate progeny resulting from self-versus male-sperm (as in G). (F) Wild type (WT) and *alh-6* hermaphrodites mated with WT males yield similar number of total progeny. (G) WT hermaphrodites mated with *alh-6;gst-4p::gfp* yield more non-GFP progeny (indicating self-fertilization) than hermaphrodites mated with wild type males harboring *gst-4p::gfp*. Statistical comparisons by unpaired t-test. *, p<0.05; **, p<0.01; ***, p<0.001; ****, p<0.0001. All studies performed in at least biological triplicate; refer to Table S1 for n for each comparison.

### *alh-6* fertility defects are sperm-specific

We noted that *alh-6* mutant hermaphrodite animals laid twice as many unfertilized oocytes as wild type animals over their reproductive-span (Figure 1D), suggesting an impairment of sperm function (Argon & Ward; McCarter, Bartlett, Dang, & Schedl; Ward & Miwa). It is notable that *alh-6* mutant hermaphrodites lay very few, if any, dead eggs (Figure 1D), suggesting that the loss of ALH-6 activity is not lethal. To determine whether the reduced brood size of *alh-6* mutants are due to a general loss of germ cells or a specific defect in oocytes or sperm, we examined the mated-fertility of these animals by mating wild type young adult (day 0-1) males to either wildtype or *alh-6* mutant virgin hermaphrodites (in wild type *C. elegans*, male sperm outcompetes hermaphrodite sperm >99% of the time (LaMunyon & Ward; Ward & Carrel) (Figure 1E). We found that the reduced fertility in *alh-6* mutant hermaphrodites is fully rescued by wild type sperm, which confirmed that oocyte quality is not impaired but rather, *alh-6* hermaphrodite sperm appears to be dysfunctional (Figure 1F).

To better assess the quality of *alh-6* mutant sperm, we compared the ability of *alh-6* mutant male sperm to compete against wild type hermaphrodite sperm (Singson, Hill, & L’Hernault). In *C. elegans* wild type animals, male sperm are larger and faster than hermaphrodite sperm, which affords a competitive advantage (LaMunyon & Ward). To differentiate between progeny resulting from mating and progeny that arise from hermaphrodite self-fertilization, we made use of male animals harboring a GFP transgene such that any cross-progeny will express GFP while progeny that arise from hermaphrodite self-sperm will not (Figure 1E). We found that wild type hermaphrodites when mated to *alh-6* mutant males have significantly more self-progeny as compared to wild type hermaphrodites mated to wild type males (Figure 1G). This finding indicates a sperm competition deficit of *alh-6* males resulting in an increased proportion of progeny derived from self-fertilization, which is uncommon after mating has occurred (Ward & Carrel). *C. elegans* hermaphrodites produce a set amount of sperm exclusively at the L4 developmental stage, before switching exclusively to oogenesis. As such, hermaphrodites eventually deplete their reservoir of sperm (Hirsh, Oppenheim, & Klass; Ward & Carrel). To assess whether *alh-6* mutant sperm are generally dysfunctional, we mated older hermaphrodites that had depleted their complement of self-sperm and found that *alh-6* mutant males are able to produce equal numbers of progeny as wild type males when the need for competition with hermaphrodite sperm is abated (Figure S3A); thus, although *alh-6* mutant sperm are impaired for competition, they remain viable for reproduction. Similarly, older sperm-depleted *alh-6* mutant hermaphrodites produced similar brood sizes when mated to young wild type or *alh-6* mutant males, which further supports a model where sperm, but not oocytes, are defective in *alh-6* mutants (Figure S3B).

### Defects in mitochondrial proline catabolism impact sperm quality

Similar to mammals, the contribution of sperm to fertility in *C. elegans* is dictated by distinct functional qualities, which include: sperm number, size, and motility (LaMunyon & Ward; Singson et al.). We next sought to define the nature of the sperm competition defect in *alh-6* mutants by measuring sperm number, size, and motility in *alh-6* mutants compared to wild type animals. One day after the on set of spermatogenesis (at the L4 larval stage of development), *alh-6* adult hermaphrodites have a reduced number of sperm in the spermatheca as compared to wild type (Figure S4A), which is correlated with the reduced self-fertility observed (Figures 1B-C). In contrast, age-matched *alh-6* mutant virgin males have similar numbers of spermatids as WT virgin males, suggesting that they have a similar rate of production (Figure 2A). We next examined sperm size in day 1 adult males and discovered that *alh-6* mutant spermatids are significantly smaller as compared to wild type (Figure 2B). To achieve motility, *C. elegans* spermatids must be activated to allow pseudopod development which requires protease activation (Ward, Hogan, & Nelson) (Figure S4B). Sperm activation can be recapitulated *in vitro* by treatment of isolated spermatids with the *Streptomyces griseus* protease Pronase (Shakes & Ward). After 30 minutes of Pronase treatment, 80% of wildtype spermatids are fully activated, while a significantly reduced population of in *alh-6* mutant spermatids mature over the same time period (Figure 2C). Interestingly, although sperm number was the same between WT and *alh-6* mutant males on the OP50 diet, sperm number was reduced in *alh-6* mutant males fed HT115 diet compared to age-matched WT males on the same diet (Figure 2D). We also noted that spermatids from *alh-6* mutant males raised on the HT115 diet were similarly defective in size and activation (Figure 2E-F). Taken together, although diet can influence sperm number, the reduction of sperm size and activation are likely contributors to the reduced fertility and competitive fitness in *alh-6* mutant males; which is independent of diet.

**Figure 2.**
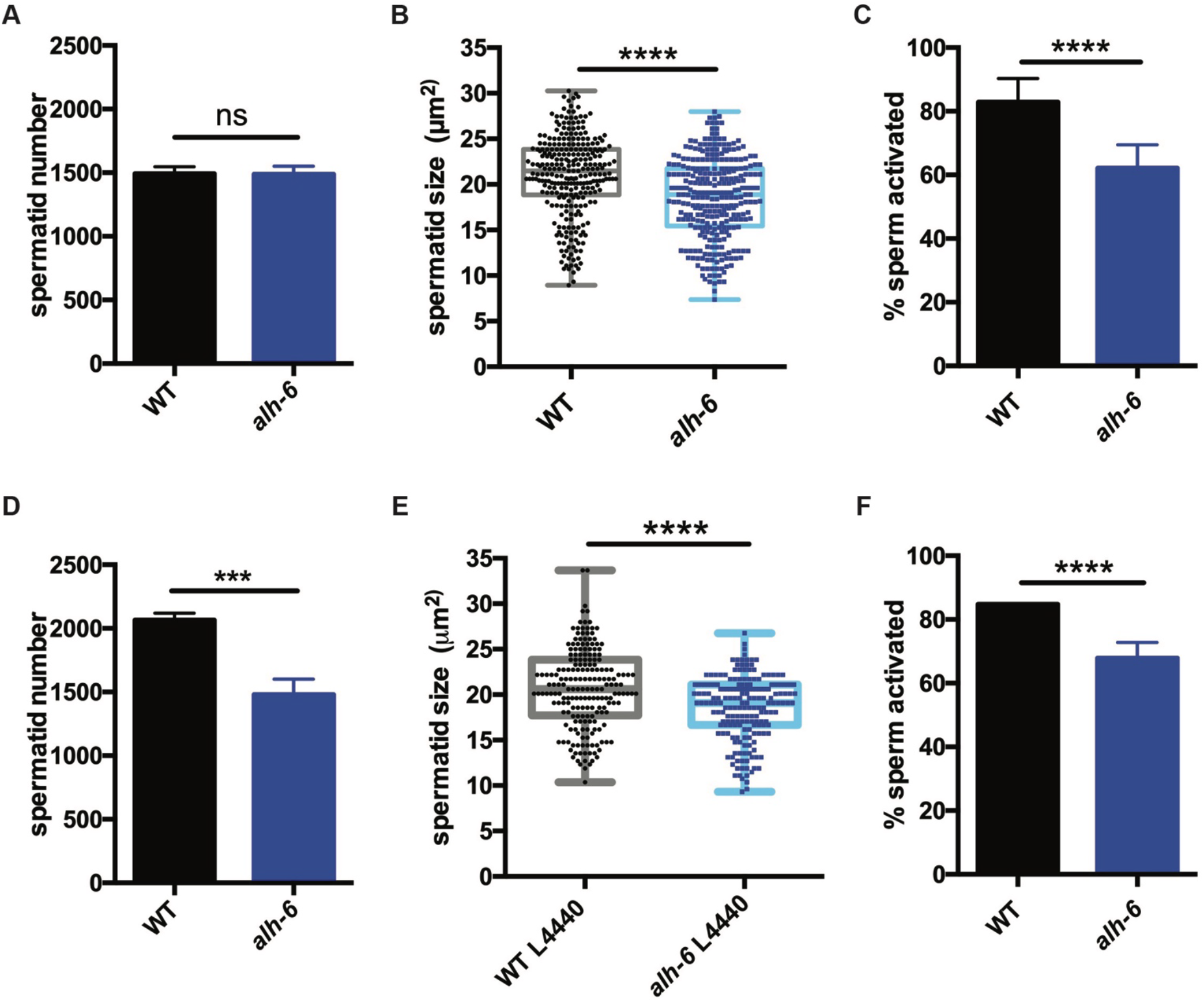
*alh-6* males have sperm defects on both OP50 and HT115 diets. (A-C) sperm phenotypes on OP50 diet. (A) Sperm quantity is similar between wild type (WT) and *alh-6* mutant day 1 adult males. (B) Spermatid size is reduced in *alh-6* mutant day 1 adult males as compared to age matched WT males. (C) Sperm activation is impaired in *alh-6* mutant day 1 adult males relative to age-matched WT males. (D-F) sperm phenotypes on HT115 diet. (D) Sperm quantity is reduced in *alh-6* mutant day 1 adult males compared to age-matched WT males. (E) Spermatid size is reduced in *alh-6* mutant day 1 adult males as compared to age matched WT males fed HT115. (F) Sperm activation is impaired in *alh-6* mutant day 1 adult males relative to age-matched WT males fed HT115. Statistical comparisons of sperm number and size by unpaired t-test and sperm activation by Fisher’s exact test. *, p<0.05; **, p<0.01; ***, p<0.001; ****, p<0.0001. All studies performed in at least biological triplicate; refer to Table S1 for n for each comparison.

### Transcriptional signatures define temporal phenotypes of *alh-6* mutant animals

We first identified *alh-6* mutant in a screen for activators of the cytoprotective transcription factor SKN-1/NRF2 using *gst-4p::gfp* as a reporter (Pang & Curran; Pang et al.). When activated, SKN-1 transcribes a variety of gene targets that collectively act to restore cellular homeostasis. However, this can come with an energetic cost with pleiotropic consequences (An & Blackwell; Blackwell, Steinbaugh, Hourihan, Ewald, & Isik; Glover-Cutter, Lin, & Blackwell; Lynn et al.; Paek et al.; Palikaras, Lionaki, & Tavernarakis; Pang & Curran; Pang et al.). *alh-6* mutants have normal development, but display progeroid phenotypes towards the end of the normal reproductive span (Pang & Curran) indicating a temporal switch in phenotypic outcomes. We reasoned that the temporally controlled phenotypes in the *alh-6* mutants could be leveraged to identify potential mechanisms by which *alh-6* loss drives cellular dysfunction. As SKN-1 is activated in *alh-6* mutants after day 2 of adulthood (Pang & Curran), we defined genes that display differentially altered expression in the L4 developmental stage, when spermatogenesis occurs, as compared to day 3 adults (post SKN-1 activation). We performed RNA-Seq analyses of worms with loss of *alh-6* and identified 1935 genes in L4 stage animals and 456 genes in day 3 adult animals that are differentially expressed (+/− Log_2_ (fold change), 0.05 FDR) (Figure S5). Notably, the gene expression changes at these two life periods had distinct transcriptional signatures (Figures 3A-B). Because the loss of *alh-6* drives compensatory changes in normal cellular metabolism, which later in life results in the activation of SKN-1, we expected to identify significant changes in both metabolic genes and SKN-1 target genes. Supporting this hypothesis, the Gene Ontology (GO) terms most enriched include oxidoreductases and metabolic enzymes in L4 stage animals (Figure 3A) and SKN-1-dependent targets such as glutathione metabolism pathway genes in day 3 adults (Figure 3B). Importantly, our transcriptomic analysis recapitulated the temporally-dependent phenotypic outcomes resulting from *alh-6* loss; genes in the pseudopodium and germ plasm GO terms class displayed reduced expression in L4 *alh-6* mutant animals (Figure 3A), which include many genes in the major sperm protein (MSP) family that comprises 15% of total protein content in *C. elegans* sperm and impact sperm function (Klass & Hirsh). In contrast, genes in the muscle-specific GO term class displayed increased expression in day 3 adults (Figure 3B), which is when SKN-1 activity is enhanced in the muscle of *alh-6* mutants (Pang et al.). Taken together, the transcriptomic analysis of *alh-6* mutants is diagnostically relevant and informative for defining drivers of organism-level phenotypic changes in animals with altered proline catabolism.

**Figure 3.**
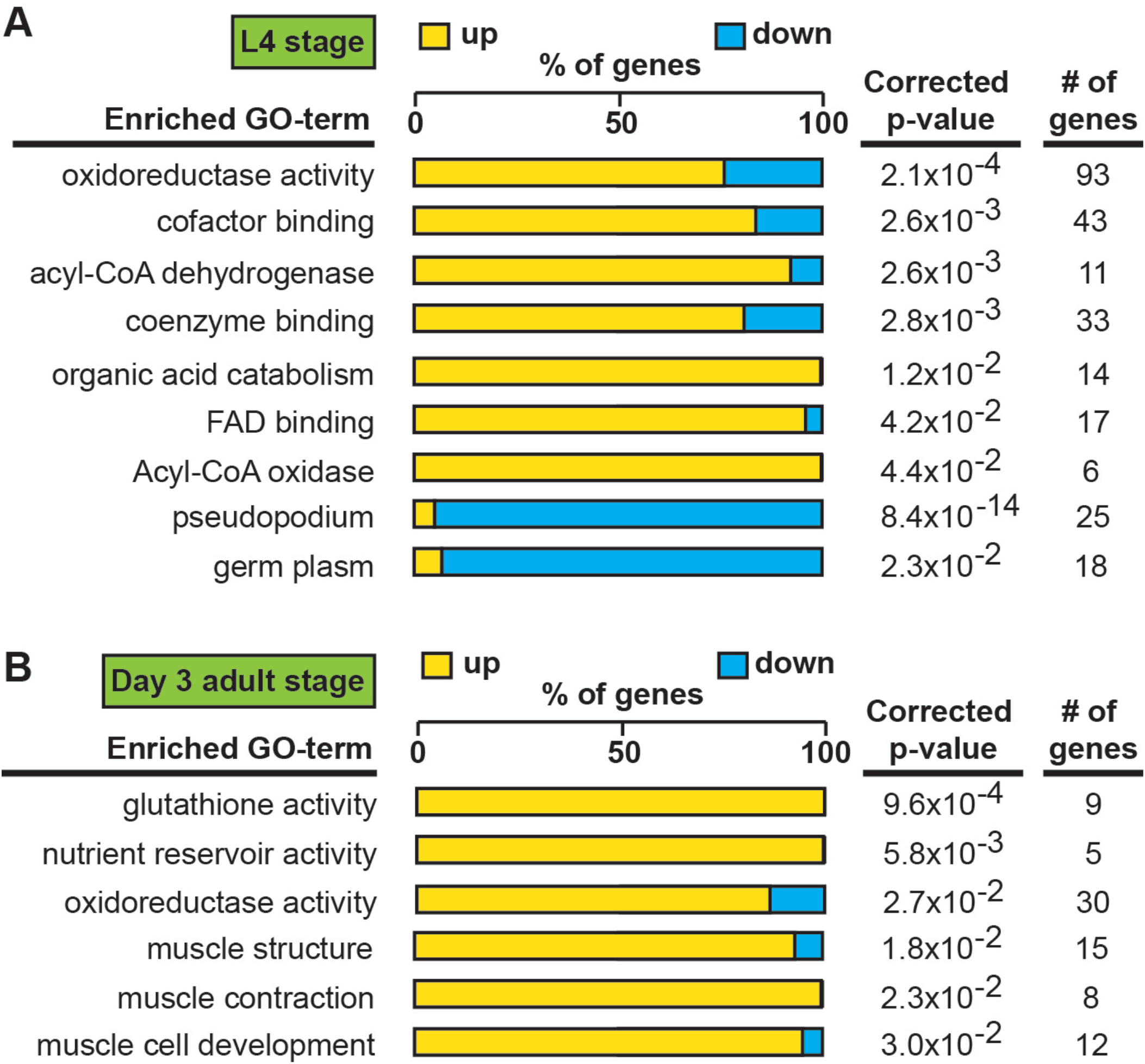
Transcriptional patterns define developmental- and adult-specific consequences to loss of *alh-6* activity. Gene Ontology (GO) term enrichment analysis of RNAseq data. (A) Transcriptional changes at L4 stage are enriched for metabolism and sperm-specific genes. (B) Transcriptional changes at day 3 adulthood are enriched for changes in glutathione activity, oxidoreductase activity, and muscle-specific genes. All studies performed in at least biological triplicate; refer to Table S1 for n for each comparison.

### FAD mediates sperm functionality and competitive fitness

The strong enrichment of genes whose protein products utilize and/or bind cofactors or co-enzymes was intriguing as the maintenance of metabolic homeostasis and the redox state of the cell requires a sophisticated balance of multiple cofactors (Figure 4A). In fact, the proline catabolism pathway utilizes multiple cofactors to generate glutamate from proline; PRDH-1 uses FAD as a co-factor to convert proline to P5C while ALH-6 utilizes the reduction of NAD+ to convert P5C to glutamate. Additionally, in the absence of ALH-6, accumulation of P5C, the toxic metabolic intermediate of proline catabolism, drives the expression of pathways to detoxify P5C (oxidoreductases, P5C reductase, etc.) (Figure 3, Figure S6A). We measured FAD and found a significant reduction in *alh-6* mutant animals fed the OP50 diet at the L4 stage (Figure 4B) and a similar reduction in animals fed HT115 bacteria at L4 stage (Figure 4C). Differences in FAD levels were unremarkable in day 3 adult animals, when spermatogenesis has long since ended (Figure S6B). Based on this finding, we predicted that restoration of FAD levels might alleviate the sperm-specific phenotypes of *alh-6* mutants. Riboflavin is a precursor of FAD (Figure 4D) and dietary supplementation of riboflavin has been shown to increase cellular FAD levels in wild-type animals (Burch, Combs, Lowry, & Padilla; Redondo, Menasche, & Le Beau). Similarly, riboflavin supplementation to the OP50 diet of *alh-6* mutants restored FAD levels to wild-type levels (Figure 4E). We found that wild type hermaphrodites mated to *alh-6* mutant males fed a riboflavin supplemented diet produced significantly more total progeny than *alh-6* males fed the standard OP50 diet (Figure S6C). Moreover, riboflavin supplementation was sufficient to partially restore male sperm size (Figure 4F) and also rescued the impaired activation (Figure 4G) of male sperm in *alh-6* mutants. Riboflavin supplementation increases sperm size in WT males, but do not change sperm activation in WT males (Figure S6D-E).

**Figure 4.**
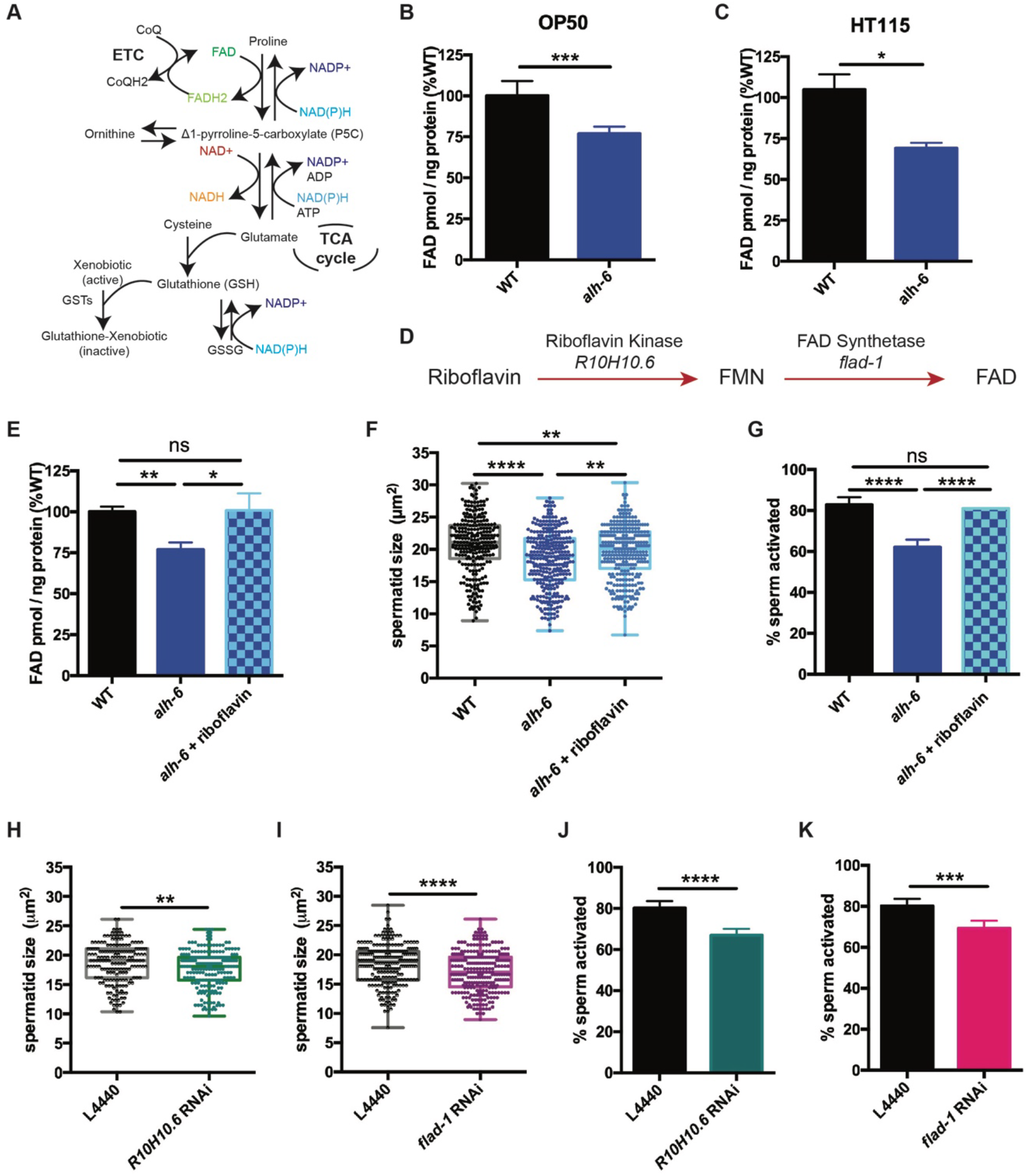
Loss of FAD homeostasis in *alh-6* mutants leads to sperm dysfunction. (A) Metabolic pathways utilize adenine dinucleotide cofactors to maintain redox balance in cells. (B-C) FAD+ levels are reduced in *alh-6* mutant animals fed OP50 (B) or HT115 (C) at the L4 developmental stage. (D) FAD biosynthetic pathway. (e-g) Dietary supplement of riboflavin restores FAD level (E), sperm size (F), and sperm activation (G) in *alh-6* mutants. (H-I) RNAi knockdown of *R10H10.*6 (H) or *flad-1* (I) in WT males reduces their sperm size compared to L4440 vector control. (J-K) RNAi knockdown of *R10H10.6* (J) or *flad-1* (K) in WT males impairs sperm activation upon Pronase treatment. Statistical comparisons of sperm size by ANOVA. Statistical comparisons of activation by fisher’s exact test with p-value cut-off adjusted by number of comparisons. *, p<0.05; **, p<0.01; ***, p<0.001; ****, p<0.0001. All studies performed in biological triplicate; refer to Table S1 for n for each comparison.

We next asked whether FAD metabolism was required for proper sperm function. FAD can be synthesized *de novo* by a two-step enzymatic reaction where riboflavin is converted to FMN by Riboflavin Kinase/R10H10.6, which is subsequently converted to FAD by FAD synthase/FLAD-1 (Figure 4D). We used RNA interference (RNAi) against *R10H10.6* or *flad-1* in wild-type male animals and measured sperm quality. Similar to *alh-6* mutant sperm, RNAi reduction of the FAD biosynthetic pathway decreased sperm size (Figure 4H, 4I, Figure S6F, S6G) and impaired sperm activation (Figure 4J, 4K, Figure S6F, S6G).

NAD+ and NADH are also central adenine dinucleotide cofactors that play critical roles in metabolism and have received recent attention as a method to combat the decline seen in biological function with age(Guarente). As such, we also measured NAD and NADH levels, but found the ratio unremarkable between wild-type and *alh-6* mutant animals (Figure S6F-H). Taken together, these data suggest that loss of *alh-6* leads to a specific decrease in cellular FAD levels and that FAD is a critical cofactor that drives proper sperm function.

### Mitochondrial dynamics regulate spermatid function

Although there is a clear and documented role for mitophagy in the clearance of paternal mitochondria post-fertilization in *C. elegans*, the role(s) for mitochondrial dynamics and turnover in sperm function prior to zygote formation are unclear. We first examined mitochondrial dynamics in wild type spermatids by staining with the fluorescent mitochondrial-specific dye JC-1 (ThermoFisher), and noted that each spermatid on average contained multiple discernable spherical mitochondria that are mostly not fused (Figures 5A, 5B, 5E). Previous studies in yeast and cultured mammalian cells have shown that when cells are exposed to mild stress, the initial response of mitochondria is to fuse in order to dilute damage (Gomes, Di Benedetto, & Scorrano; Rambold, Kostelecky, Elia, & Lippincott-Schwartz; Tondera et al.). Indeed *alh-6* mutant spermatids had mitochondria that were more interconnected (Figures 5C-E) as compared to wild type spermatids, which suggests that *alh-6* sperm are under metabolic stress. The increase in fused mitochondria in spermatids was also present in animals fed the HT115 diet, which further supports a diet-independent role for *alh-6* in the germline (Figure S7A).

**Figure 5.**
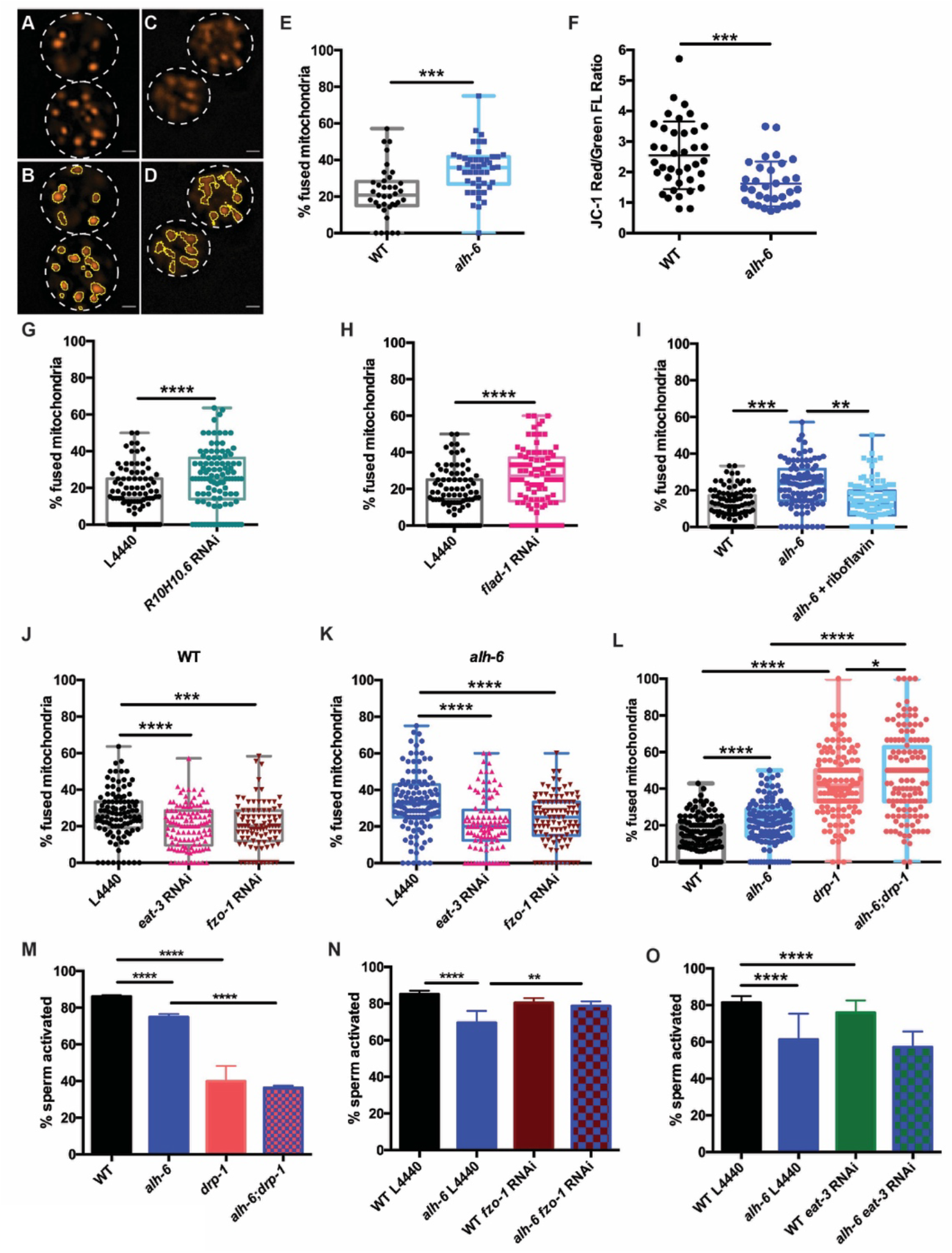
Mitochondrial dynamics drive sperm quality. (A-E) JC-1 dye stained mitochondria of WT (A-B), *alh-6* mutant (C-D); (B and D) are ImageJ detection of JC-1 stained sperm mitochondria area which are quantified in (E). (F) Mitochondria in *alh-6* mutant spermatids have reduced JC-1 red/green fluorescence ratio, indicating mitochondria depolarization. (G-H) RNAi knockdown of FAD biosynthetic pathway genes, *R10H10.6* (G) or *flad-*1 (H) increases mitochondrial fusion in WT spermatids. (I) Dietary supplement of FAD precursor riboflavin restores mitochondrial fusion in *alh-6* spermatids to WT level. (J-K) *eat-3* or *fzo-1* RNAi decreases mitochondrial fusion in both WT (J) and *alh-6* (K) mutant spermatids. (L) *drp-1* mutation increases mitochondrial fusion in spermatids. (M) *drp-1* mutation significantly impairs sperm activation in both WT and *alh-6* mutant spermatids. (N) *fzo-1* RNAi restores sperm activation in *alh-6* mutant. (O) *eat-3* RNAi reduces sperm activation in WT males but not *alh-6* males. Statistical comparisons of JC-1 Red/Green FL ratio by unpaired t-test. Statistical comparisons of mitochondria fusion by ANOVA. Statistical comparisons of sperm activation by Fisher’s exact test with p-value cut-off adjusted by number of comparisons. *, p<0.05; **, p<0.01; ***, p<0.001; ****, p<0.0001. All studies performed in at least biological triplicate; refer to Table S1 for n for each comparison.

The JC-1 dye accumulates in mitochondria in a membrane potential-dependent manner, and as concentration exceeds threshold, its fluorescence switches from green to red emission; thus, a higher red-to-green fluorescence ratio is indicative of healthier mitochondria. *alh-6* mutant spermatids have reduced red:green JC-1 fluorescence that indicates a lower mitochondrial membrane potential, and an accumulation of unhealthy mitochondria (Figure 5F).

A connection between mitochondrial dynamics (fusion and fission) and FAD homeostasis has not been previously described. To understand this, we perturbed FAD biosynthesis pathway and then examined mitochondrial connectivity in spermatids. We first reduced FAD biosynthesis with RNAi targeting *R10H10.6* or *flad-1*, which resulted in more connected mitochondria that resembles the increased fusion in *alh-6* mutant spermatids that are under metabolic stress (Figure 5G-H, Figure S6A). In addition, increasing FAD levels by dietary supplementation of riboflavin, restored mitochondria in spermatids of *alh-6* animals to more wild-type-like distributions (Figure 5I), but did not change mitochondrial morphology in WT male spermatids (Figure S7B). Thus, the reduction of FAD in *alh-6* mutants, alters mitochondrial dynamics to a more fused and less punctate state. Therefore, the homeostatic control of FAD level is critical to maintain proper mitochondrial dynamics in sperm.

The role of mitochondrial dynamics in the maturation of sperm has not been studied; however recent work has revealed that the mitochondrial fusion and fission machinery are important for the elimination of paternal mitochondria post-fertilization (Wang et al.). FZO-1 is required for proper fusion of the mitochondrial outer membrane while EAT-3/OPA1 regulates inner membrane fusion. In opposition to the activities of FZO-1 and EAT-3, DRP-1 is required for mitochondrial fission (Lima et al.; Smirnova, Griparic, Shurland, & van der Bliek). The balance of this fusion and fission machinery in the upkeep of mitochondrial homeostasis allows cells to respond to changes in metabolic needs and external stress (Shaw & Nunnari; van der Bliek, Sedensky, & Morgan). RNAi of *fzo-1* or *eat-3* reduced mitochondrial fusion in wild-type male sperm (Figure 5J and Figure S7C) and suppressed the enhanced fusion observed in *alh-6* mutant spermatid mitochondria (Figure 5K); indicating mitochondrial fusion of both membranes is active in spermatids with impaired proline catabolism. We next examined spermatids from *drp-1* mutant animals and observed a greater level of mitochondrial fusion as compared to wild type and *alh-6* mutant spermatids (Figure 5L). We observed a synergistic level of mitochondrial fusion in spermatids derived from *alh-6; drp-1* double mutants. This finding is consistent with previous studies in yeast which reveal that defects in fusion can be compensated for by changes in the rates of fission and vice versa (Shaw & Nunnari; van der Bliek et al.). In support of our model where mitochondrial dynamics act as a major driver of the sperm-specific defects in *alh-6* mutants, we discovered that loss of *drp-1*, which results in increased mitochondrial fusion (like that observed in *alh-6* mutants), also reduces sperm activation (Figure 5M). Moreover, reducing *fzo-1* or *eat-3* does not alter activation in wild type sperm, while *fzo-1* but not *eat-3* RNAi restores activation in *alh-6* sperm (Figure 5N-O and Figure S7C), suggesting increased fusion mediated predominantly by *fzo-1* in *alh-6* sperm mitochondria is impairing proper function. We noted that *alh-6* mutant animals have an increased expression of *fzo-1* transcripts that is suggestive of a retrograde signaling response from the mitochondria (Figure S7D). Taken together, these data support a model where loss of mitochondrial proline catabolism induces mitochondrial stress, activating mitochondrial fusion, in order to dilute damage to preserve functional mitochondria at the cost of sperm function. These data also reveal a functional role for mitochondrial fusion and fission in spermatid development and sperm function.

### *alh-6* and FAD are cell autonomous regulators of sperm function

Signaling between germ and somatic cells can alter function in each cell type (J. Berman & C. Kenyon; J. R. Berman & C. Kenyon; Curtis, O’Connor, & DiStefano; Ghazi, Henis-Korenblit, & Kenyon; Greenwald; Hsin & Kenyon; Libina, Berman, & Kenyon; Lin, Hsin, Libina, & Kenyon). In light of the differences between somatic and germline phenotypes observed in *alh-6* mutant animals, we performed germline specific RNAi targeting *alh-6* to deduce whether the sperm defects observed were cell autonomous. Germline specific RNAi of *alh-6* in wild-type males was not sufficient to alter sperm size (Figure 6A), but did result in diminished sperm activation (Figure 6B,) and increased mitochondrial fusion in sperm (Figure 6C). We conclude that the effects of loss of *alh-6* on sperm function are cell autonomous because germline specific RNAi could phenocopy the sperm defects observed in whole animal loss of *alh-6*, (while the effect of *alh-6* on sperm size is non-cell autonomous and requires somatic input (Figure S8A-D). Since FAD functions in a variety of essential cellular processes, we next asked if proper sperm function required FAD homeostasis in germ cells. Similarly, we reduced *R10H10.6* or *flad-1* only in the germline, which phenocopies germline knockdown *of alh-6* on sperm size (Figure 6D,G), sperm activation (Figure 6E, H), and mitochondrial fusion in sperm (Figure 6F, I), as observed in whole animal RNAi of *flad-1* or *R10H10.6* (Figure 4J-K and Figure 5G-H).These results suggest that FAD functions similarly to *alh-6* in cell autonomously regulating sperm function (activation and mitochondrial dynamics), while affecting sperm size in a cell non-autonomous manner (Figure 4H-I). Lastly, we restored wild-type *alh-6* expression, only in the germline, in *alh-6* mutant animals, which restored sperm size in one of the two transgenic lines (Figure 6J), activation (Figure 6K) and mitochondrial dynamics (Figure 6L), as compared to non-transgenic siblings. Taken together these data identify the importance of proline catabolism and FAD homeostasis in germ cells to maintain proper sperm function. In conclusion, our studies define mitochondrial proline catabolism as a critical metabolic pathway for male reproductive health.

**Figure 6.**
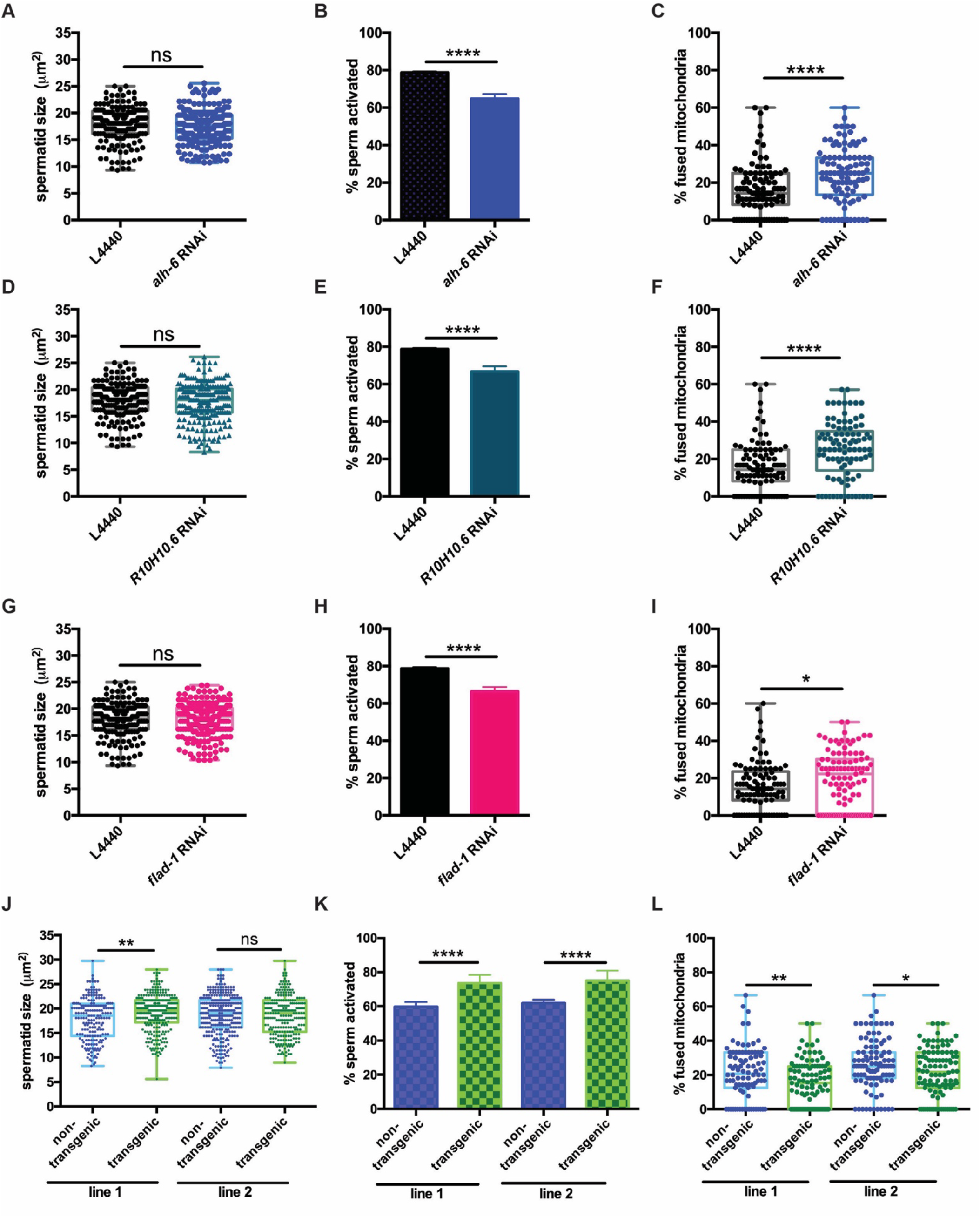
*alh-6* and FAD function cell autonomously in the germline to regulate sperm function. (A-C) Germline-specific RNAi of *alh-6* does not change sperm size (A), but does impair sperm activation (B) and increases mitochondrial fusion in sperm (C). Similarly, (D-I) germline-specific RNAi of *R10H10.6* and *flad-1* do not change sperm size (D,G), impair sperm activation (E,H), and increase mitochondrial fusion in sperm (F,I). (K-L) Germline-specific rescue of WT *alh-6* in *alh-6* mutant male animals increases sperm size (J) and restores activation (K) and mitochondrial dynamics (L). Statistical comparisons of sperm size and mitochondrial fusion in spermatids by unpaired t-test. Statistical comparisons of sperm activation by Fisher’s exact test with p-value cut-off adjusted by number of comparisons. *, p<0.05; **, p<0.01; ***, p<0.001; ****, p<0.0001. All studies performed in biological triplicate; refer to Table S1 for n for each comparison.

## DISCUSSION

Here we investigate the effects of disrupting mitochondrial proline catabolism through the loss of the mitochondrial enzyme gene *alh-6* and the resulting changes in FAD homeostasis, mitochondrial dynamics, and male fertility (Figure 7). We found that *alh-6* mutants show a reduction in brood size that is sexually dimorphic; defects in sperm function but not oocytes contribute to reduced hermaphrodite fertility. As societal factors continue to push individuals to wait longer to have children, the increase in paternal age is inversely correlated with proper sperm function and can give rise to fertility issues. Consequently, it is incumbent on future studies to elucidate how restoring and maintaining functional amino acid catabolism during aging in order to promote reproductive success.

**Figure 7.**
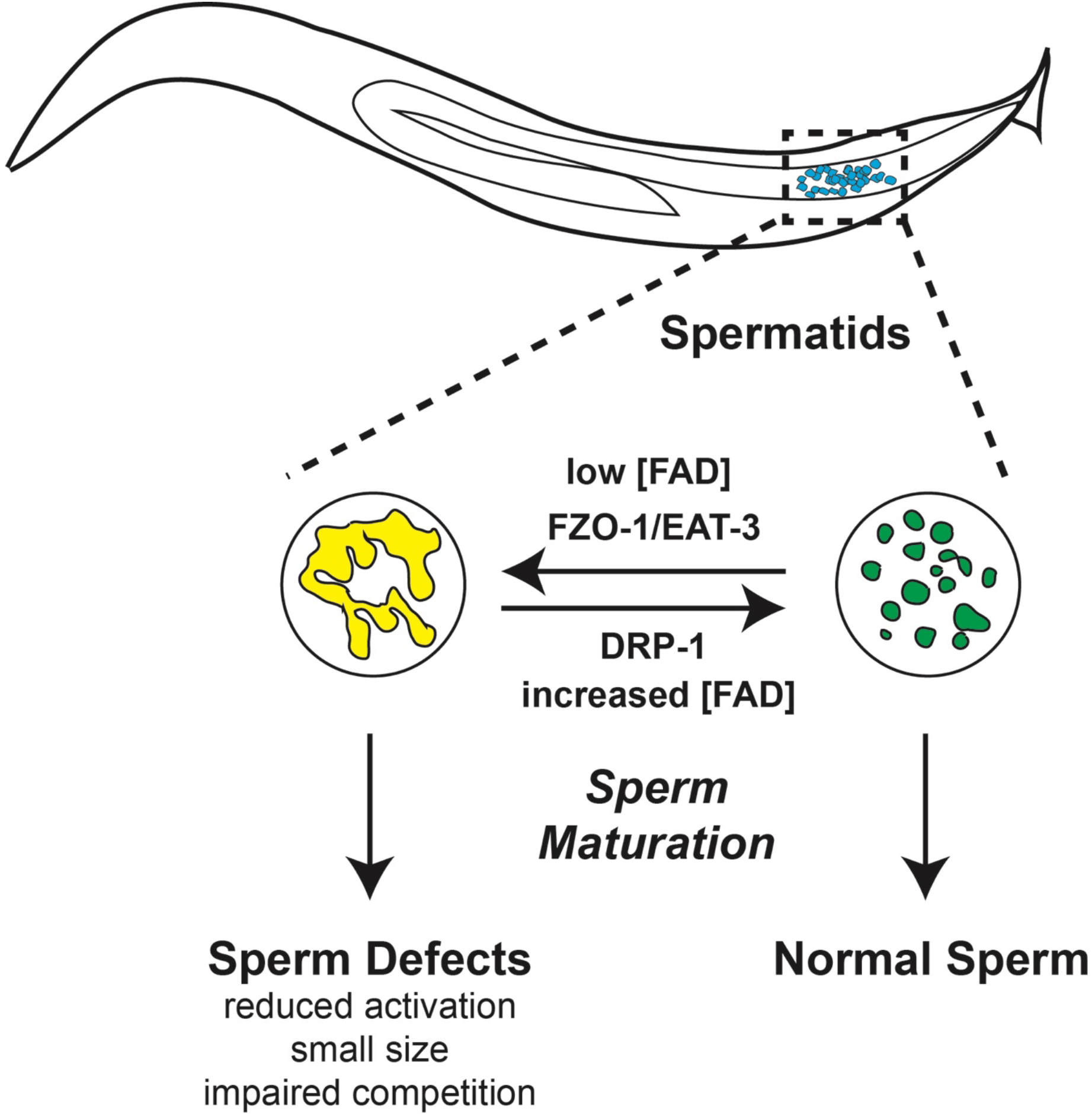
Model of *alh-6* and FAD mediated male reproductive senescence.

Although *C. elegans* is a well-established organism for studying aging and reproduction, with several studies describing hermaphrodite reproductive senescence, many questions regarding the basis of male reproductive decline remain unanswered. Decades of work have shown that exposure to pollution, toxins, xenobiotics, and other ROS-inducing compounds can prematurely drive the loss of sperm function across species (Agarwal, Virk, Ong, & du Plessis; Cocuzza, Sikka, Athayde, & Agarwal; Wagner, Cheng, & Ko), but the impact that normal cellular metabolism plays on sperm function and the identification of specific molecules that can mediate sperm quality are not well-defined. In this study we characterized a new role for mitochondrial proline catabolism and FAD homeostasis in the maintenance of proper sperm function. Perturbation of this pathway, through mutation of *alh-6/ALDH4A1*, causes metabolic stress. Consequently, this perturbation leads to reduction of cellular FAD level and increases mitochondrial fusion in spermatids, which results in impaired sperm function and premature reproductive senescence.

Mutation in proline dehydrogenase (*PRODH*) in humans results in hyperprolinemia type I (HPI), while mutation in delta-1-pyrroline-5-carboxylate dehydrogenase (*ALDH4A1/P5CDH*) results in hyperprolinemia type II (HPII). This study reveals that in *C. elegans*, proline catabolism impacts several functional qualities of male sperm. Loss of proline catabolism results in smaller sperm with impaired activation, two qualities that directly impact competitive advantage. As such, proline biosynthesis, catabolism, and steady state concentrations must be tightly regulated, and the importance of proline in cellular homeostasis may help explain the transcriptional responses measured in animals with dysfunctional *alh-6*. Our data support a cell autonomous role for proline catabolism in sperm. However, although whole animal RNAi of *alh-6* closely phenocopies the *alh-6* mutant including reduced spermatid size, germline specific RNAi of *alh-6* did not significantly reduce the size of spermatids; perhaps suggesting a partial role for ALH-6 in somatic tissues for spermatid development, which is in line with recent studies in *C. elegans* describing soma to germline signaling in sperm activation (Chavez, Snow, Smith, & Stanfield). Intriguingly, the impact of loss of *alh-6* is mostly independent of diet source, unlike the somatic phenotypes which are diet-dependent (Pang & Curran). The exception is sperm number in *alh-6* mutant animals on the HT115 diet, which appears to be diet-dependent (Figure 2D). WT males have more spermatids when fed the HT115 diet, as compared to WT animals fed OP50 diet, while *alh-6* mutants have the same number of spermatids on both diets.

Our previous work defined the age-dependent decline in function of somatic tissues, particularly muscle in animals lacking functional ALH-6 (Pang & Curran; Pang et al.), which does not manifest until day 3 of adulthood. Our current study reveals that although somatic phenotypes in *alh-6* mutants are observed post-developmentally, the germline, or more specifically spermatids, are sensitive to loss of *alh-6* much earlier in development (phenotypes assayed at L4 or Day 1 of adulthood). Reproductive senescence is a field of growing significance as the number of couples that choose to delay having children increases. Importantly, although *alh-6* mutant sperm are impaired for competition, they remain viable for reproduction. This is similar to recent study on *comp-1*, a mutation which results in context-dependent competition deficit in *C. elegans* sperm (Hansen, Chavez, & Stanfield).

Recent studies have focused on the role of NAD+ metabolism in cellular health, while the impact of FAD has received less attention. FAD levels are diminished in *alh-6* animals specifically at the L4 stage when spermatogenesis is occurring. Riboflavin (Vitamin B_2_) is a precursor to the FAD and FMN cofactors that are needed for metabolic reactions in order to maintain proper cellular function, like proline catabolism and mitochondrial oxidative phosphorylation. Despite its importance, humans, like *C. elegans*, lack a riboflavin biosynthetic pathway and therefore require riboflavin from exogenous sources (Powers). Insufficient intake can lead to impairment of flavin homeostasis, which is associated with cancer, cardiovascular diseases, anemia, neurological disorders, impaired fetal development, etc. (Powers). Our study suggests that riboflavin and FAD play critical roles in reproduction, specifically in germ cell development, as loss of FAD biosynthesis or loss of *alh-6* specifically in the germline recapitulates the sperm defects observed in whole animal knockdown or *alh-6* mutation. Importantly, these sperm-specific defects can be corrected by dietary supplementation of vitamin B_2_, which in light of the exceptional conservation of mitochondrial homeostatic pathways, suggest the nutraceutical role vitamin B_2_ could play in sperm health across species.

Our study also demonstrates that spermatids lacking *alh-6* have increased mitochondrial fusion; a perturbation at the mitochondrial organelle structure-level that contributes to the sperm-specific phenotypes observed. In addition to prior work showing *fzo-1/MFN1/MFN2* and *drp-1*/*DRP-1* to be important for mitochondrial elimination post-fertilization (Wang et al.), our work reveals that mitochondrial fission and fusion machinery are present and active in spermatids and that perturbation of these dynamics can affect sperm maturation and competitive fitness. Future work to define how *alh-6* spermatids use mitophagy, which can clear damaged mitochondria, will be of interest. In conclusion, our work identifies proline metabolism as a major metabolic pathway that can impact sperm maturation and male reproductive success. Moreover, these studies identify specific interventions to reverse the redox imbalance, cofactor depletion, and altered mitochondria dynamics, all of which play a part in sperm dysfunction resulting from proline metabolism defects.

## ACKNOWLEDGEMENTS

We thank N. Mih, K. Han, and L. Thomas for technical assistance and H. Dalton, A. Hammerquist, N. Stuhr, W, Escorcia, and J. Nhan for critical reading of the manuscript. Some strains were provided by the CGC, which is funded by the NIH Office of Research Infrastructure Programs (P40 OD010440). This work was funded by the NIH (R01GM109028, R01AG058610, to S.P.C., T32AG000037 to D.L.R., and T32GM118289 to C.D.T), and the American Federation of Aging Research (C-A.Y. and S.P.C).

## AUTHOR CONTRIBUTIONS

S.P.C. designed the study; C-A.Y. performed the majority of all experiments with assistance from D.L.R., C.D.T., S.P., and S.P.C.; C-A.Y. and S.P.C. analyzed data. S.P.C. wrote the manuscript and C-A.Y. and S.P.C. revised the manuscript. All authors discussed the results and commented on the final version of the manuscript.

## COMPETING INTERESTS

The authors declare no competing interests.

## DATA AVAILABILITY

All relevant data are available from the authors. RNA-Seq data are deposited in GEO database (GSE121920).

## METHODS

### *C. elegans* strains and maintenance

*C. elegans* were cultured using standard techniques at 20°C. The following strains were used: wild type (WT) N2 Bristol, SPC321[*alh-6(lax105)*], SP326[*alh-6p::alh-6::gfp*], SPC447[laxEx025 - *pie-1p::alh-6*], SPC455[laxEx033 - *pie-1p::alh-6*], SPC473[laxEx051 - *pie-1p::alh-6*], CL2166[*gst4-p::gfp*], SPC223[*alh-6(lax105);gst-4p::gfp*], DCL569[mkcSi13(sun-1p::rde-1::sun-1 3’UTR + unc-119(+)) II; rde-1(mkc36) V], and CU6372[*drp-1(tm1108)*]. Double and triple mutants were generated by standard genetic techniques. *E. coli* strains used were as follows: B Strain OP50(Brenner) and HT115(DE3) [F^−^ mcrA mcrB IN(rrnD-rrnE)1 lambda^−^ rnc14::Tn10 λ(DE3)](Timmons, Court, & Fire). For dietary supplement assays, riboflavin was added to the NGM plate mix to final concentration 2.5mM.

### RNAi-based experiments

RNAi experiments were done using HT115-based RNAi (Timmons et al.), which yielded similar results as OP50 RNAi *E. coli* B strain as described in (Dalton & Curran). All strains were adapted to diets for at least three generations and strains were never allowed to starve. All RNAi clones were sequenced prior to use and RNAi knockdown efficiency measured. RNAi cultures were seeded on IPTG plates and allowed to induce overnight prior to dropping eggs on them for experiments.

### Microscopy

Zeiss Axio Imager and ZEN software were used to acquire all images used in this study. For GFP reporter strains, worms were mounted in M9 with 10mM levamisole and imaged with DIC and GFP filters. For sperm number, assay samples were imaged with DIC and DAPI filters in z-stacks. For sperm size and activation assays, dissected sperm samples were imaged at 100x with DIC filter on two different focal planes for each field to ensure accuracy. For sperm mitochondria assays, dissected sperm samples were imaged at 100x with DIC, GFP, and RFP filters in z-stacks to assess overall mitochondria content within each spermatid.

### Fertility assay

Worms were treated with alkaline hypochlorite and eggs were allowed to hatch overnight. The next day, synchronized L1 larvae were dropped on NGM plates seeded with either OP50 or HT115. 48 hours later, at least ten L4 hermaphrodites for each genotype were singled onto individual plates and moved every 12 hours until egg laying ceased. Progeny were counted 48 hours after the singled hermaphrodite was moved to a different plate. Plates were counted twice for accuracy.

### Mated reproductive assay

Males were synchronized by egg laying, picked as L4 larvae for use as young adults for mating experiments. Singled L4 stage hermaphrodites were each put on a plate with 30ul of OP50 seeded in the center together with three virgin adult males. 24 hours post-mating, males were removed, and each hermaphrodite was moved to a new plate every 24 hours until egg laying ceased. Progeny were counted 48 hours after the hermaphrodite was moved from the plate. For sperm competition assay, progeny with GFP fluorescence were counted from the cohort. Plates were counted twice for accuracy.

### Cofactor Measurements

Worms were treated with alkaline hypochlorite and eggs were allowed to hatch overnight. The next day, synchronized L1s were dropped on NGM plates with or without supplement seeded with 25X concentrated OP50. FAD levels are measured following directions in FAD Colorimetric/Fluorometric Assay Kit (K357) from BioVision. NAD/NADH levels are measured following directions in NAD/NADH Quantification Colorimetric Kit (K337).

### Sperm Number Assay

Worms were treated with alkaline hypochlorite and eggs were allowed to hatch overnight. The next day, synchronized L1s were dropped on NGM plates with the indicated food source. At 48 hours (L4 developmental stage) males were isolated to new plates. 72 hours post-drop, day 1 adult virgin male animals were washed 3x with 1xPBST, fixed with 40% 2-propanol, and stained with DAPI for 2 hrs. Samples were washed for 30min with PBST, mounted with vectashield mounting medium, and covered with coverslip to image. Spermatids in the seminal vesicle were counted through all planes in z-stack.

### Sperm Size Assay

Males were isolated at L4 stage 24 hours before assay. For each strain, five day 1 adult males were dissected in 35μL pH 7.8 SM buffer (50mM HEPES, 50mM NaCl, 25mM KCl, 5mM CaCl_2_, 1mM MgSO_4_, 10mM dextrose) to release spermatids, which were immediately imaged.

### Sperm Activation with Pronase

Males were isolated at L4 stage 24 hours before assay. For each strain, five day 1 adult males were dissected in 35μL pH 7.8 SM buffer (50mM HEPES, 50mM NaCl, 25mM KCl, 5mM CaCl_2_, 1mM MgSO_4_, 1mg/ml BSA) supplemented with 200μg/mL Pronase® (Millipore Sigma) to release spermatids. Another 25ul of the same solution was added and the spermatids were incubated at RT for 30 min for activation to occur.

### Sperm Mitochondria Staining

Males were isolated at L4 stage 24 hours before assay. For each strain, five day 1 adult males were dissected in 35μL pH 7.8 SM buffer (50mM HEPES, 50mM NaCl, 25mM KCl, 5mM CaCl_2_, 1mM MgSO_4_, 1mg/ml BSA) with JC-1(Thermo Fisher Scientific T3168) added to 10μM final concentration. Another 25ul of the same solution was added and the spermatids were incubated at RT for 10 min. The slide was washed three times with 100ul SM buffer before imaging.

### RNA-sequencing

Worms were egg prepped and eggs were allowed to hatch overnight. The next day, synchronized L1s were dropped on NGM plates seeded with 25X concentrated OP50. 48 and 120 hours post drop, L4 animals and day 3 adult animals, respectively, were washed 3 times with M9 and frozen in TRI Reagent at −80°C. Animals were homogenized and RNA extraction was performed following the protocol in Zymo Direct-zol RNA Isolation Kit. RNA samples were sequenced and analyzed by Novogene.

### Statistical analysis

Data are presented as mean ± SEM. Comparisons and significance were analyzed in Graphpad Prism 7. Comparisons between two groups were done using Student’s Test. Comparisons between more than two groups were done using ANOVA. For sperm activation assays, Fisher’s Exact Test was used and p-values are adjusted for multiple comparisons. *p<0.05 **p<0.01 *** p<0.001 ****<0.0001.

## SUPPLEMENTAL FIGURES

**Figure S1.**
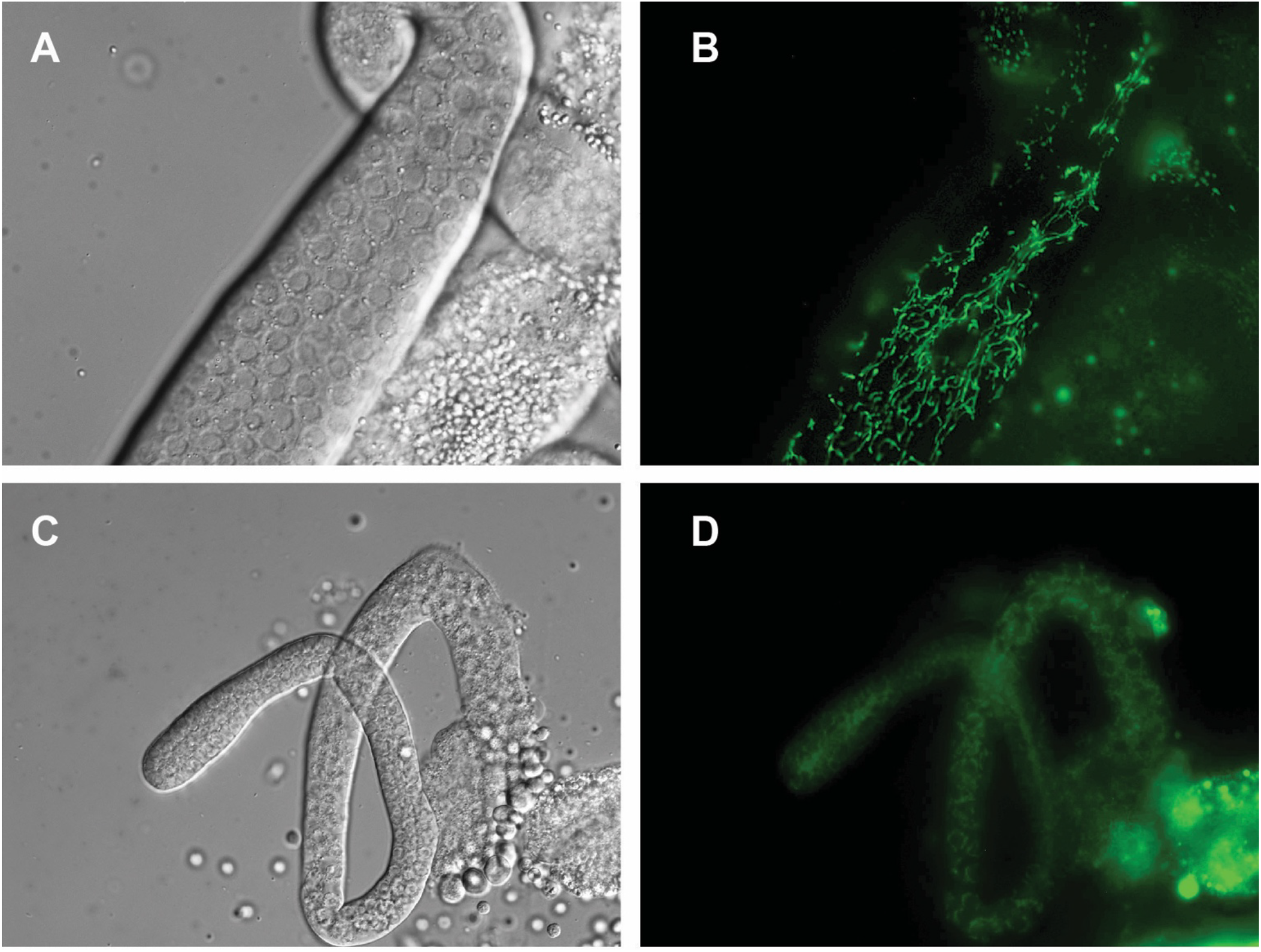
Mitochondrial localization of ALH-6 in the germline. UV integrated *alh-6::gfp* strain under its endogenous promoter reveal expression of ALH-6 in hermaphrodite (A-B) and male (C-D) germline. a and c are DIC images while b and b are GFP images.

**Figure S2.**
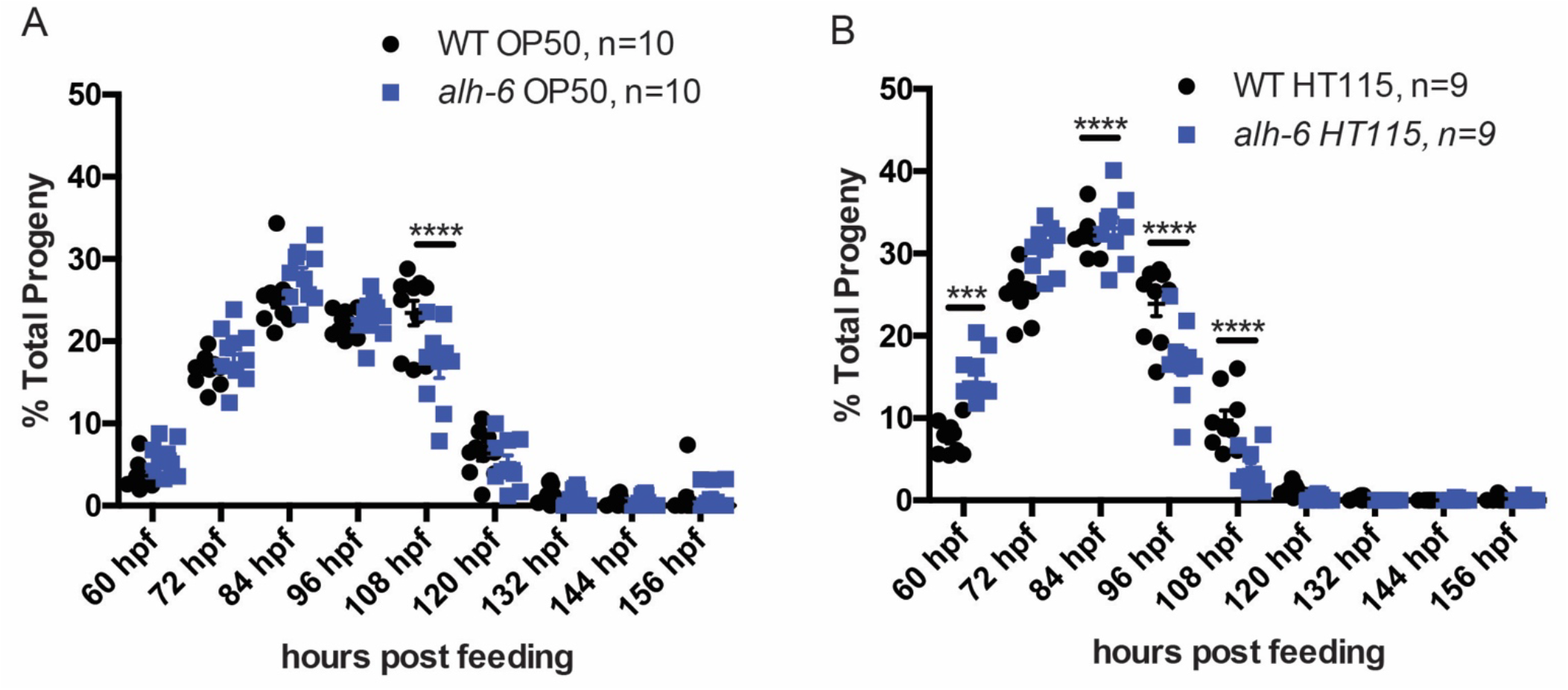
*alh-6* hermaphrodite reproductive span is similar to wild type (WT) on different diets. Progeny output time-courses are plotted as % total progeny for each time point. WT and *alh-6* mutant have similar output on OP50 (A) and HT115 (B). Significance indicate differences in progeny output at a particular time point done by multiple t-tests. *, p<0.05; **, p<0.01; ***, p<0.001; ****, p<0.0001. All studies performed in biological triplicate; refer to Table S1 for n for each comparison.

**Figure S3.**
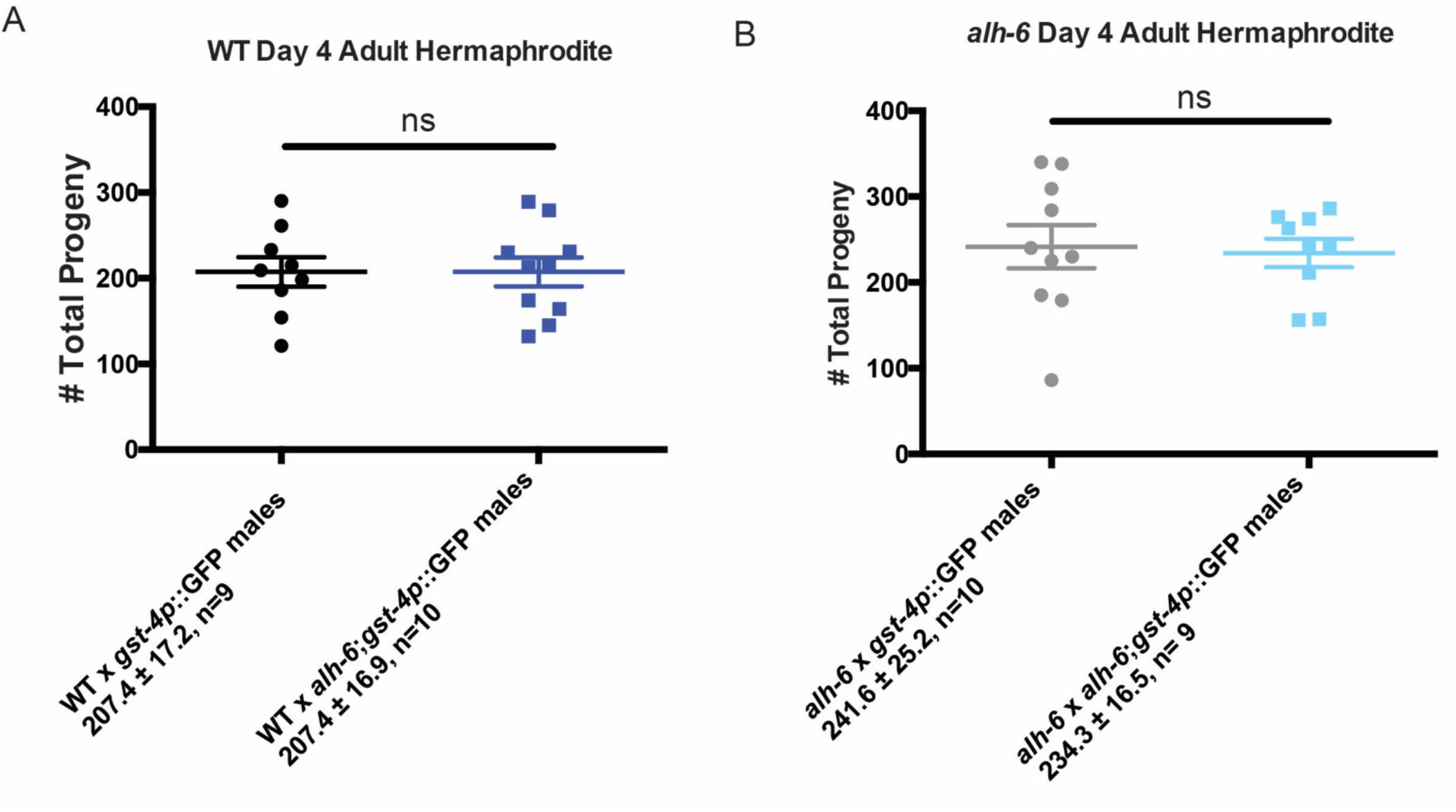
*alh-6* fertility defects are sperm-specific. (A) Day 4 adult WT hermaphrodites mated to either *gst-4p::gfp* or *alh-6;gst-4p::gfp* males yield similar total brood size. (B) Day 4 adult *alh-6* hermaphrodites mated to either *gst-4p::gfp* or *alh-6;gst-4p::gfp* males yield similar total brood size. Comparisons made with unpaired t-test. *, p<0.05; **, p<0.01; ***, p<0.001; ****, p<0.0001. All studies performed in biological triplicate; refer to Table S1 for n for each comparison.

**Figure S4.**
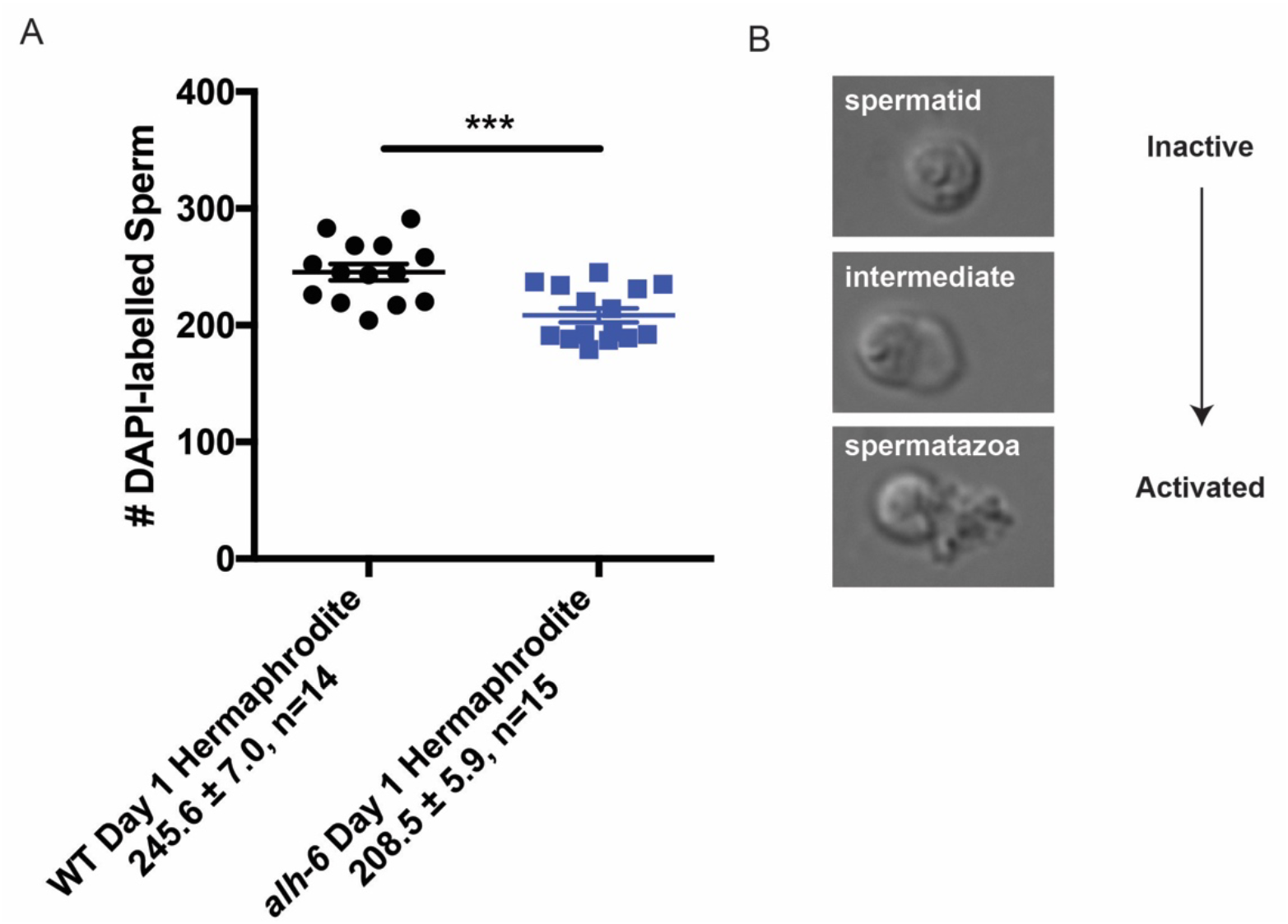
Sperm defects in *alh-6* mutants. (A) *alh-6* hermaphrodites have reduced sperm number as day 1 adults. (B) Spermiogenesis stages. Spermatozoa with fully formed pseudopods are considered activated.

**Figure S5.**
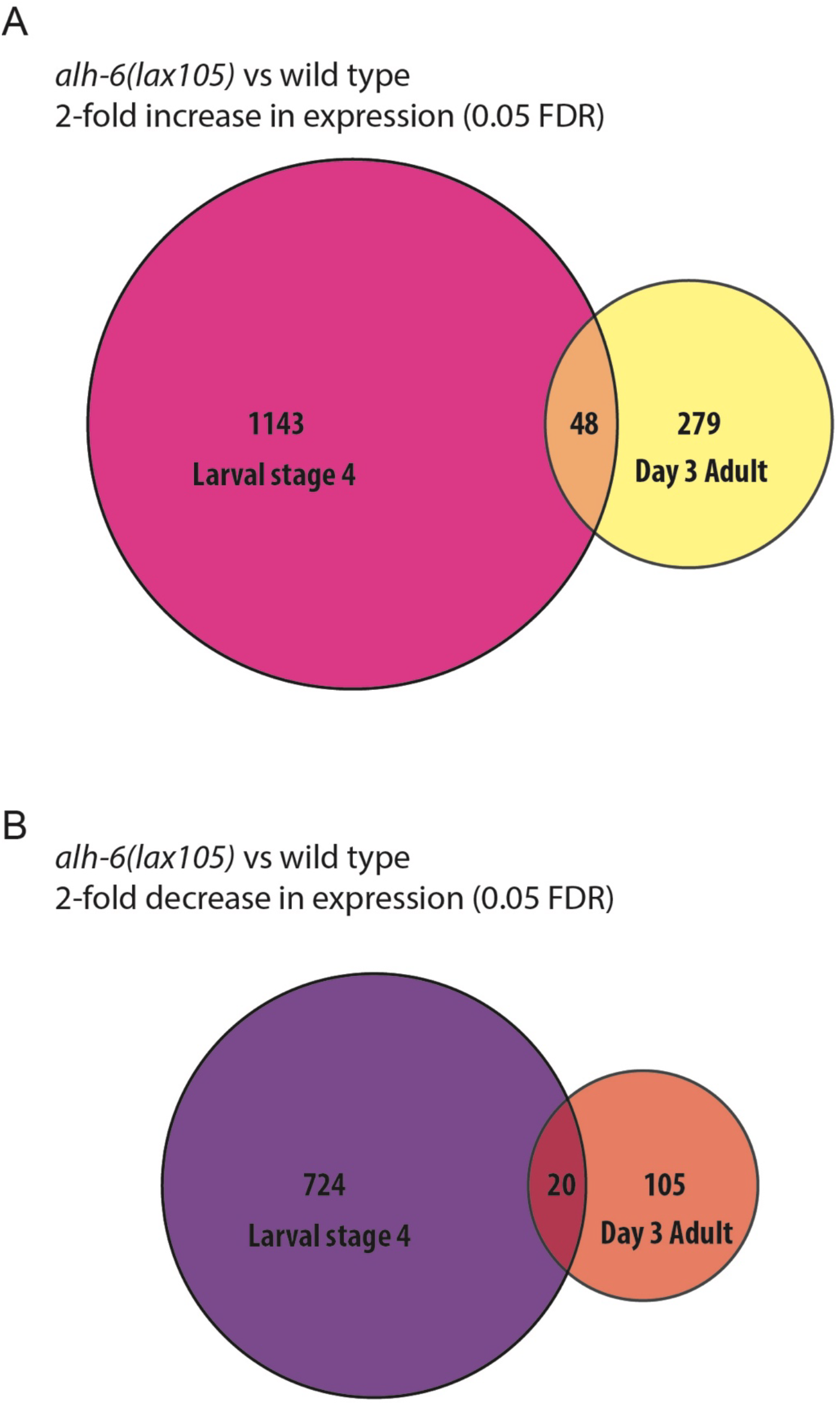
RNA-Sequencing data of WT and *alh-6* hermaphrodites at L4 and day 3 adulthood. (A) Number of genes that are significantly upregulated in *alh-6(lax105)* compared to WT at L4 and Day 3 adult stages. (B) Number of genes that are significantly downregulated in *alh-6(lax105)* compared to WT at L4 and Day 3 adult stages. FDR = 0.05. All studies performed in biological triplicate; refer to Table S1 for n for each comparison.

**Figure S6.**
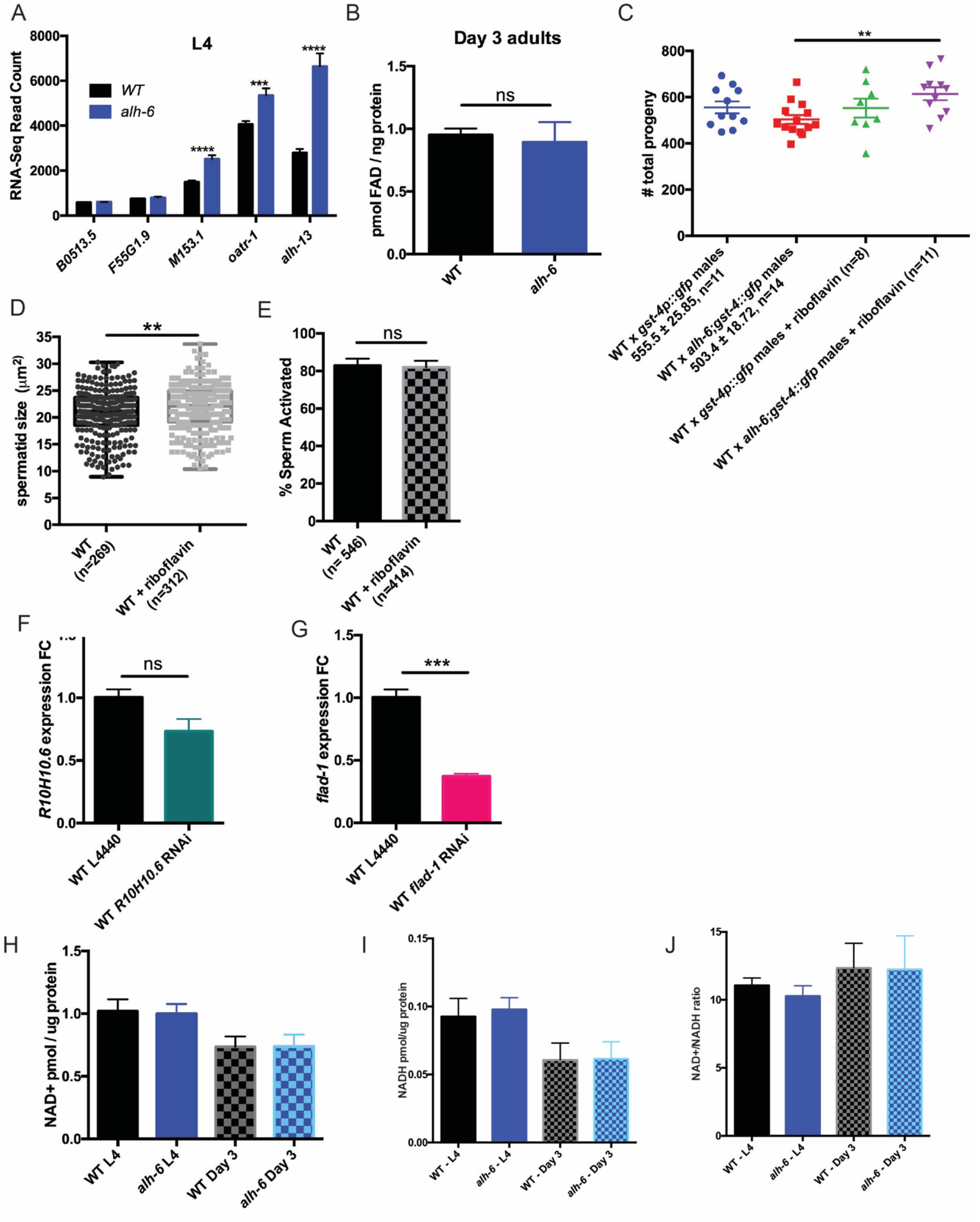
Adenine nucleotide cofactor homeostasis is disrupted in *alh-6* mutants. (A) *alh-6* mutants display increased expression of metabolic enzymes to reduce P5C levels. (B) FAD levels are unchanged between WT and *alh-6* mutant animals fed OP50 at day 3 adulthood. (C) WT hermaphrodites mated to *alh-6;gst-4p::gfp* males fed OP50 supplemented with 2.5mM riboflavin results in increase in total brood size compared to non-supplemented *alh-6;gst-4p::gfp* males. (D) Dietary riboflavin supplement increased sperm size of WT males. (E) WT males fed OP50 diet supplemented with riboflavin have similar % spermatid activated upon Pronase treatment as those without riboflavin supplement. (F) *flad-1* expression is reduced by whole animal RNAi via RT-PCR verification. (G) *R10H10.6* expression is modestly reduced by whole animal RNAi. (H) NAD levels are unchanged between WT and *alh-6* animals at L4 and day 3 adulthood (I) NADH level is unchanged between aged matched WT and *alh-6* hermaphrodites at both L4 and Day 3 adulthood. (J) NAD+/NADH level is unchanged between aged matched WT and *alh-6* hermaphrodites at both L4 and Day 3 adulthood. *, p<0.05; **, p<0.01; ***, p<0.001; ****, p<0.0001. All studies performed in biological triplicate; refer to Table S1 for n for each comparison.

**Figure S7.**
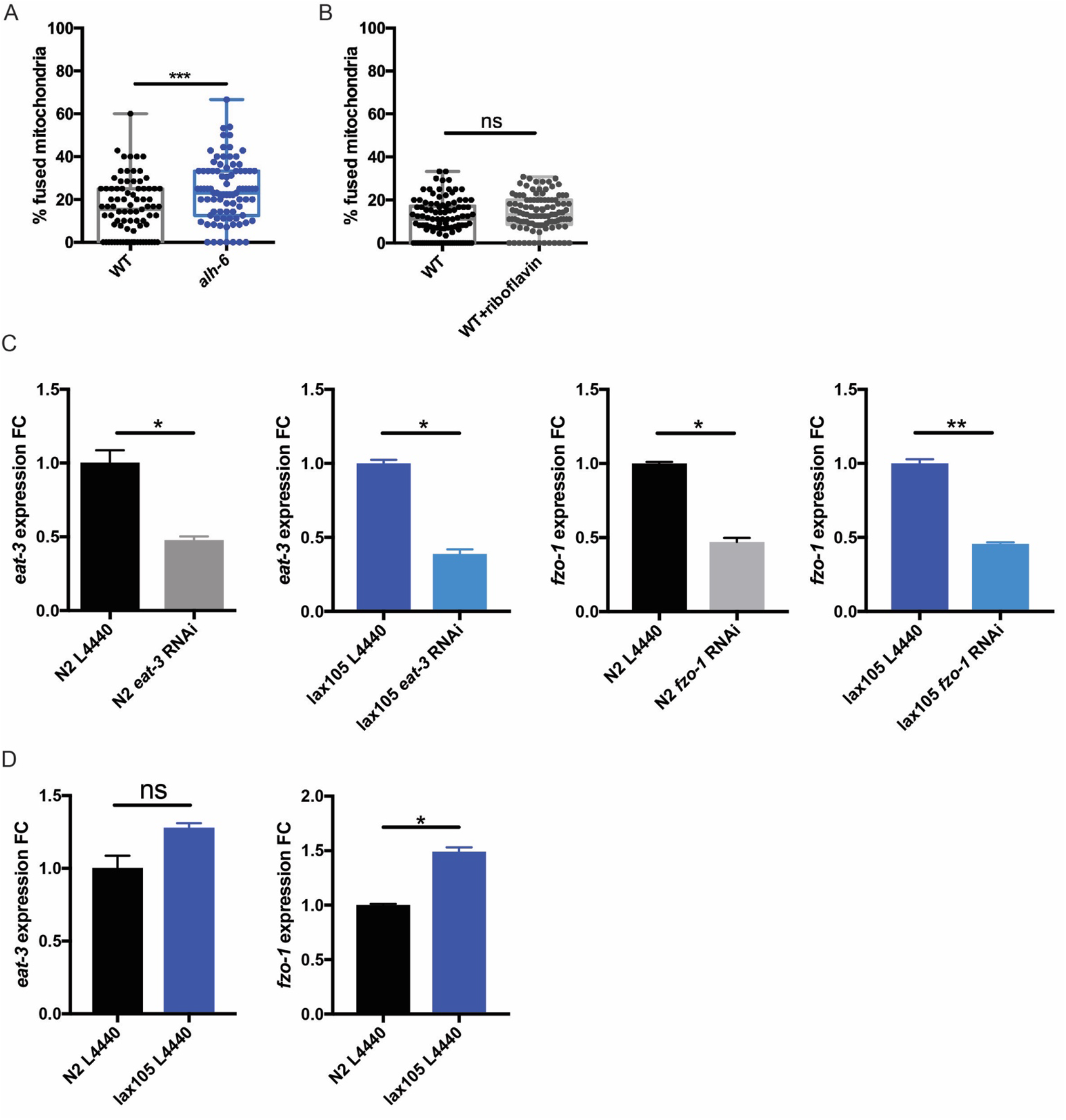
*fzo-1* is involved in mitochondrial dynamics aberration in *alh-6* spermatids. (A) *alh-6* males fed HT115 diet still have increased mitochondrial fusion in spermatids compared to age-matched WT males. (B) Riboflavin supplementation did not alter mitochondria in spermatids of WT males. (C) *eat-3* and *fzo-1* RNAi knockdown in WT and *alh-6* mutants are verified using RT-PCR. (D) *fzo-1* expression is increased in *alh-6* mutants, while *eat-3* expression is not significantly increased.

**Figure S8.**
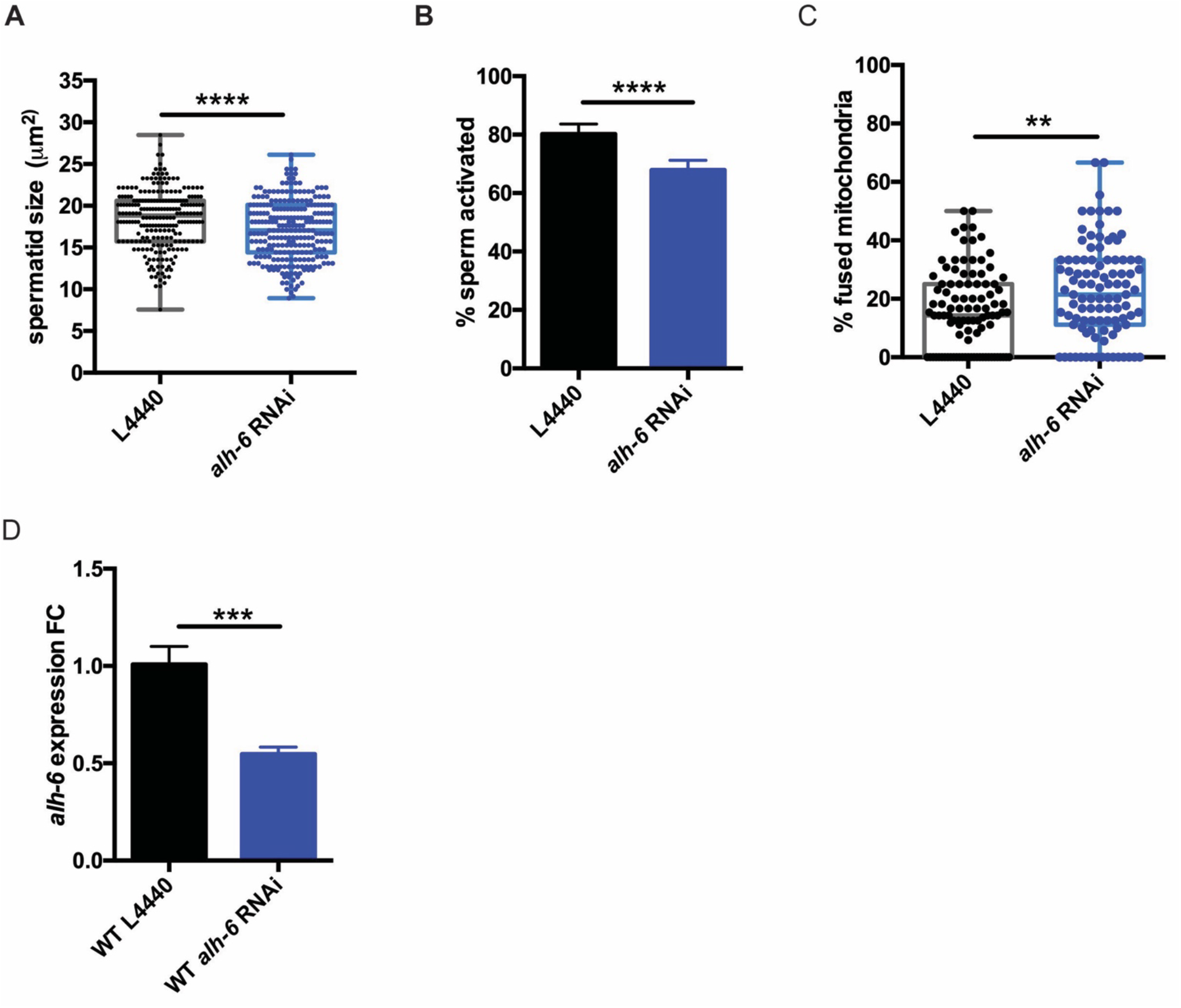
Germline expression of WT *alh-6* is sufficient to rescue sperm defect. (A-C) Whole animal RNAi of *alh-6* reduces sperm size (A), impairs activation (B), and increases mitofusion in spermatids (C). (D) *alh-6* expression is reduced by whole animal RNAi via RT-PCR verification.

